# Extensive Loss and Gain of Conserved Non-Coding Elements during Early Teleost Evolution

**DOI:** 10.1101/2023.05.23.541954

**Authors:** Elisavet Iliopoulou, Vasileios Papadogiannis, Costas S. Tsigenopoulos, Tereza Manousaki

## Abstract

Conserved Non-coding Elements (CNE) in vertebrates are enriched around transcription factor loci associated with development. However, loss and rapid divergence of CNEs has been reported in teleost fish, albeit taking only few genomes into consideration. Taking advantage of the recent increase in high-quality teleost genomes, we focus on studying the evolution of teleost CNEs, carrying out targeted genomic alignments and comparisons within the teleost phylogeny to detect CNEs and reconstruct the ancestral teleost CNE repertoire. This teleost-centric approach confirms previous observations of extensive vertebrate CNE loss early in teleost evolution, but also reveals massive CNE gain in the teleost stem-group over 300 million years ago. Using synteny-based association to link CNEs to their putatively regulated target genes, we show the most teleost gained CNEs are found in the vicinity of orthologous loci involved in transcriptional regulation and embryonic development that are also associated with CNEs in other vertebrates. Moreover, teleost and vertebrate CNEs share a highly similar motif and transcription factor binding site vocabulary. We suggest that early teleost CNE gains reflect a restructuring of the ancestral CNE repertoire through both extreme divergence and *de novo* emergence. Finally, we support newly identified pan-teleost CNEs have potential for accurate resolution of teleost phylogenetic placements in par with coding sequences, unlike ancestral only elements shared with spotted gar. This work provides new insight into CNE evolution with great value for follow-up work on phylogenomics, comparative genomics and the study of gene regulation evolution in teleosts.

## Introduction

During embryonic development, the interaction of a network of Transcription Factors (TFs) with a variety of cis-regulatory elements found mainly in the non-coding part of the genome drives the spatial and temporal regulation of gene expression with great precision^1^. In vertebrates, regulatory elements can be close to target genes (< 50kbp) or even embedded in intronic areas, but many cis-regulatory modules are located from hundreds of kbp to more than 1 Mb away^2^. Among the non-coding genome, a large number of vertebrate CNEs have maintained greater than 70% sequence identity for over 400 million years, exhibiting even greater average conservation than protein-coding genes^3^. Many of these CNEs act as enhancers for neighbouring genes associated with evolutionarily conserved functions, including important developmental processes^4^. While most vertebrate CNEs are found in a single copy in the genome, some CNE duplications have also been identified in paralogous loci produced by the two rounds (2R event) of Whole Genome Duplication (WGD) that happened early in vertebrate evolution, suggested to have contributed to the diversification of the subphylum ^5^.

While jawed vertebrates share thousands of CNEs, a few hundred of these elements, including duplicate CNEs, are even conserved with lamprey, a member of the basally splitting jawless vertebrates^6^. Therefore, the evolution of a set of ancestral vertebrate CNEs predates the split of jawless and jawed vertebrates, possibly dating to the first WGD shared by both lineages, with additional CNEs established after the second WGD of jawed vertebrates. Within vertebrates, teleost fish form the most species-rich monophyletic group^7^. It is hypothesized that one of the main drivers of their widespread diversification is the teleost-specific (3R event) WGD^8^. Because of 3R, teleost genomes have increased genetic diversity, carrying additional paralogous genes. Unexpectedly, in contrast to gene coding regions, earlier reports have suggested extensive loss and rapid divergence of ancestral vertebrate CNEs following the 3R WGD event^8,9^, though this has been based on few model species and comparisons to other vertebrates. To date, the drivers of this massive loss of CNEs after 3R have been unexplored, while the conservation of non-coding elements within the teleost clade has remained uncharacterized.

Previous studies on teleost CNE evolution had mainly focused on the fate of ancestral vertebrate CNEs in teleost fish^10^. Therefore, a comprehensive catalogue of non-coding elements conserved within the teleost group is lacking and the way the CNE landscape responded to the 3R WGD remains unresolved. Here, we focus on identifying and studying teleost CNEs through a teleost-centric search, with the main focus of investigating the evolution of CNEs following the 3R WGD. For this purpose, we carry out targeted genome alignments of key focal species and use these data to detect CNEs across the teleost phylogeny. Drawing information of CNE presence and absence in different teleost groups, we reconstruct the ancestral teleost CNE repertoire and test the potential of these ancestral teleost CNEs as markers for resolving phylogenetic teleost placements. We then assess sequence and synteny conservation of teleost CNEs and use an orthology-guided synteny-based approach to associate CNE gains and losses with candidate conserved gene targets with putative regulatory links to studied CNEs. Finally, we use motif discovery and enrichment analysis to compare the transcription factor binding site composition of gained and ancestral CNEs. This work represents the first CNE analysis on a large number of teleost genomes and as such it provides unique insight for understanding the evolution of the conserved non-coding genome of teleosts.

## Materials and Methods

All analyses were carried out on the Zorbas HPC infrastructure of IMBBC – HCMR ^11^.

To identify teleost CNEs we used the identification pipeline described below and presented in Supplementary Figure 1. Briefly, we first selected 24 teleost genomes and the spotted gar genome from the Ensembl database^12^ (Supplementary table 1), aiming for high contiguity (N50 > 8Mb), high completeness (BUSCO score of > 95%) and broad sampling across the teleost phylogeny. We then used pairwise whole genome alignments from two selected pairs of focal reference species for *de novo* CNE identification and used these starting CNE sets to identify ancestral teleost elements, as well as CNE gains and losses across the phylogeny.

**Figure 1.**
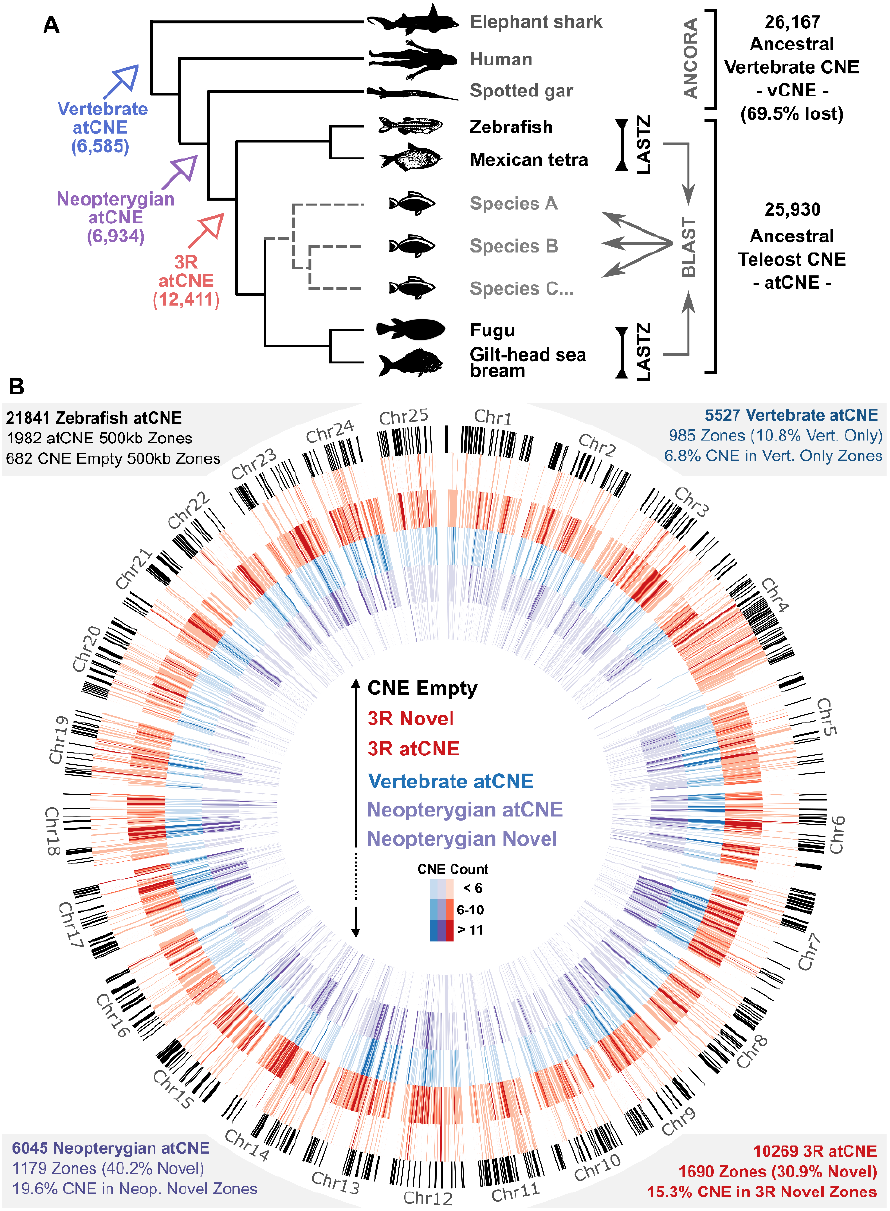
Teleost ancestral CNE identification and distribution. **A**. CNE sets from pairwise whole genome alignments in two focal reference genome pairs (LASTZ) were searched in 20 other teleost genomes via BLAST, identifying 25,930 atCNEs that were present in the teleost common ancestor. Comparison to non-teleost vertebrate genomes and ancestral vertebrate CNEs from the ANCORA database revealed widespread loss of vertebrate atCNEs and gain of novel atCNEs in Neopterygians and Teleosts (3R). **B**. Distribution and count of 21,841 atCNEs over 1,982 500kb-long zones in the zebrafish genome. From largest (outermost) to smallest (innermost) circle radius: 1) Genomic regions with no atCNE presence (black). 2) Novel zones with 3R atCNE presence (red) and absence of older (neopterygian or vertebrate) atCNE. 3) Zones with 3R atCNE presence (red). 4) Zones with vertebrate atCNE presence (blue). 5) Zones with Neopterygian atCNE presence (purple). 6) Novel zones with Neopterygian atCNE presence (purple) and absence of older (Vertebrate) atCNE.

## CNE Identification pipeline

### Pairwise whole-genome alignment

We performed pairwise reciprocal whole-genome alignments with LASTZ v1.04.03 ^13^ for two focal reference pairs of genomes downloaded from the Ensembl database^14^, *Danio rerio* (Zebrafish - DanRer11) VS *Astyanax mexicanus* (Mexican tetra - Astyanax_mexicanus-2.0) and *Takifugu rubripes* (Fugu - fTakRub1.2) VS *Sparus aurata* (Gilthead sea bream - fSpaAur1.2). These pairs were selected as representatives of the two most basally splitting clades of teleost phylogeny (206 - 252 MYA ^15^), to allow for inference of ancestral CNE sets through comparisons across the phylogeny. All genomes were downloaded hard masked, *i.e*., interspersed repeats and low complexity regions were detected with the RepeatMasker tool^16^ and were replaced with N. To conduct the alignments, we split input genomes in 10Mb windows with 100kb overlaps for Zebrafish and Fugu and non-overlapping 10Mb windows for Mexican tetra and gilthead seabream. Two rounds of pairwise whole-genome alignments were run for each pair as discussed by Hiller *et al* 2013 ^17^. In the first round, we used the following more sensitive LASTZ parameterization recommended for distantly related species (> 100 Mya): M=50 E=30 H=2000 K=2200 L=6000 O=400 T=1 Y=3400 Q=HoxD55.q ^18^.Subsequently, a second round of alignment was performed after masking aligned regions from the first alignment round, to facilitate the identification of conserved elements not found in the initial alignment. During the second round, the following less sensitive LASTZ parameters were used: K=1500, L=2300, M=0 and W=5 ^17^.

### Chaining - Netting

Aligned regions found by the two rounds of pairwise whole-genome alignment were chained using the CNEr v1.8.3 Bioconductor wrapper functions^18^ inside an R environment (using utilities found in UCSC Browser ^19^). During chaining, matching sufficiently close initial alignments are joined into bigger alignment chains, with the longest fragments further connected to form netted alignments in net Axt format (Supplementary Figure 1).

### CNE Identification in reference focal species

Alignment chain nets from the two focal reference pairs (Zebrafish VS Mexican tetra and Fugu VS Gilthead seabream) were used for *de novo* CNE identification using Zebrafish and Fugu as reference genomes. Exonic regions were filtered out of alignments, using gene annotation information from the Ensembl database (*Takifugu_rubripes.fTakRub1.2.104, GCF_900880675.1_fSpaAur1.1_genomic, Astyanax_mexicanus-2.0.104, Danio_rerio.GRCz11.104*). The CNEr v1.28.0 Bioconductor package^18^ was used for both filtering and CNE detection. Conserved regions were selected using the following parameters: 1) a maximum number of 4 hits per element (cutoffs = 4), i.e., how many times we expect to see an element, 2) a minimum identity of 70% for aligned regions (Identities = 70pc), *i.e*., the minimum percentage of matches in a single alignment and 3) a sliding window size of 100bp for detection (windows = 100).

### CNE search in other teleost species

Using the *de novo* identified CNE datasets from Zebrafish (*zCNEs*) and Fugu (*fCNEs*), we searched 20 other teleost fish genomes pre-masked for repeats via BLAST v2.10.0+ ^20^ for zCNE and fCNE presence / absence (evalue <= 1e^-06^, word size=6, max target seqs=1, max hsps=1). This comparative dataset was used as the basis for studying CNE conservation, gain and loss, as well as CNE based phylogenomic analyses.

### CNE-based phylogenomic analysis

For CNE-based phylogenomic reconstructions, we used single copy CNEs shared by all species in the phylogeny, including or excluding spotted gar (*Lepisosteus oculatus*) as an outgroup species belonging to the Holostei infraclass of ray-finned bony fish, which diverged before the 3R WGD ^21^. To extract CNE sequences for multiple alignments from different teleost or spotted gar, we used BLAST derived coordinates extended by 50bp in either side to maximize alignable sequence capture (Supplementary Figure 2). Multiple sequence alignments were carried out using MAFFT v7.407 ^22^ and CNE alignments were collated with custom bash scripting to produce a single supermatrix of all CNE from across all species in each phylogeny. TrimAl v1.4.rev15 ^23^ was used for alignment trimming with a gap threshold of 50% of (gt 0.5). RAxML-NG v.1.0.2 ^24^ was used for CNE-based tree construction using 20 starting trees (10 parsimony + 10 random) and 100 bootstraps, with TVM+I+G4 selected as the best-fit model with ModelTest-NG^25^. For visualization purposes, FigTree v1.4.4 ^26,27^ was used. Gene-based reconstruction was carried out with the same pipeline, using single copy orthologous proteins for all species included predicted by OrthoFinder2 v2.5.4 ^28^ from the proteomes of these species. Protein sequences were aligned with MAFFT v7.407, collated to a supermatrix with subsequent trimming using TrimAL v1.4.rev15 (gt 0.5). Tree construction was performed with RAxML-NG v.1.0.2 using 20 starting trees (10 parsimony + 10 random) and 100 bootstraps, with JTT+I+G4+F selected as the best-fit model with ModelTest-NG.

### Identifying the vertebrate CNE set

For inference of ancestral vertebrate CNE losses in teleosts, CNE datasets for *Homo sapiens* (Human), *Callorhinchus milii* (Elephant shark) and *Lepisosteus oculatus* (Spotted gar) (using Human as reference) were downloaded from the ANCORA database and were re-filtered for coding regions (*H. sapiens* gene predictions from RefSeq, Genbank, CCDS, UniProt and the UCSC KnownGene track), selecting sequences with >70% identity over a window of 100 bases. These ancestral vertebrate CNE were then searched in the 24 teleost genomes in the study using BLAST (evalue=1e-6, word size=6, max target seqs=1, max hsps=1).

### Ancestral teleost CNEs Identification and Categorization

We based our ancestral CNE identification protocol on comparisons between the two earliest splitting teleost clades in our phylogeny hereby defined as: Clade 1 (highlighted in orange in Figure 2A) spanning from *Siluriformes* represented by *P. hypophthalmus* to *Cypriniformes* represented by *D. rerio* and Clade 2 (highlighted in blue in Figure 2A) spanning from *Cyprinodontiformes* represented by *K. marmoratus* to *Tetraodontiformes represented by M.mola*. To infer the ancestral teleost CNE set (from now on abbreviated as atCNEs), we selected zCNEs with presence in any clade two species and fCNEs not overlapping zCNEs with presence in any clade one species.

We then searched for atCNEs presence in spotted gar or other vertebrates (ghost shark, western clawed frog, chicken, green anole and human) via BLAST (e-value=1e-6, word size=6, max target seqs=1, max hsps=1) and allocated atCNEs to the following categories based on their inferred time of origin: 1) Elements identified in any vertebrate other than spotted gar were categorized as ‘vertebrate atCNEs’, inferred to be ancestral to ray-finned fish, 2) Elements present in spotted gar and teleosts and absent from all other vertebrates were categorized as ‘neopterygian atCNEs’, inferred to have been gained in the common neopterygian ancestor with spotted gar, 3) Elements present only in teleosts were categorized as ‘3R atCNEs’, inferred to have been gained in the teleost common ancestor.

**Figure 2.**
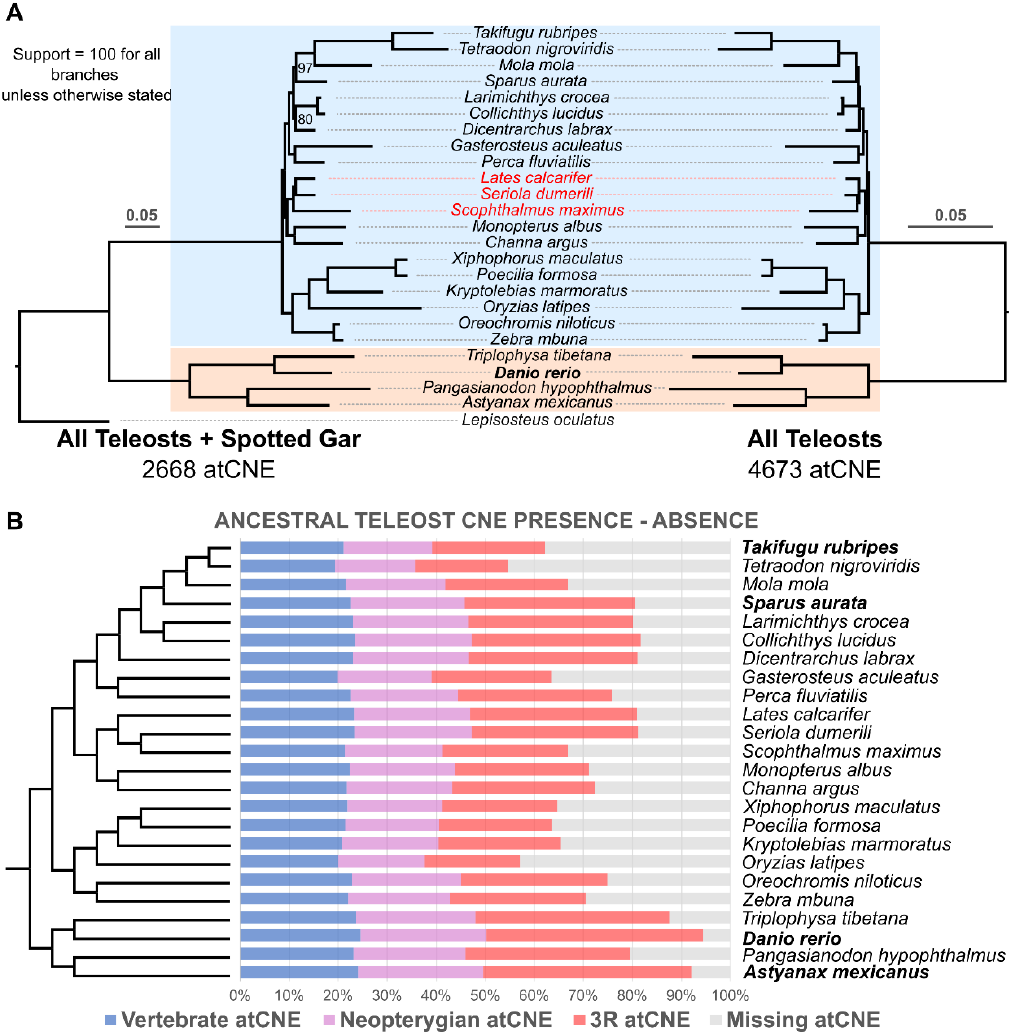
Ancestral CNE losses across the teleost phylogeny. **A**. Phylogenetic reconstruction of the relationships of the 24 teleost species included in the study, using single copy universal atCNEs, including (left tree – 2668 atCNEs) or excluding (right tree – 4673 atCNEs) spotted gar. The two basally diverged clades used to infer ancestral teleost CNEs are highlighted with orange (clade 1) and blue (clade 2). Branches with species highlighted in red show topological discrepancies in the spotted gar shared atCNE phylogeny (left). **B**. Presence / absence of ancestral atCNE categories of different origin (Vertebrate in blue, Neopterygian in purple, 3R in red) across the teleost phylogeny as a percentage of total identified atCNEs. The cladogram is constructed based on phylogenetic reconstructions from Figure 2A.

CNE distribution and localization over zebrafish chromosomes was plotted using Circos ^29^(Figure 1B) and Rideogram ^30^ (Figure 6).

### CNE Sequence and Synteny Conservation Analyses

To calculate CNE sequence conservation scores in three representative genomes from different parts of the phylogeny (*D. rerio, O. niloticus* and *T. rubripes)*, we carried out multiple whole genome alignments for all 25 species in the study with CACTUS (v 2.2.1) 31, applied phyloFit ^32^ for phylogenetic model fitting (using the REV substitution model) for each chromosome for each species and then used phyloP ^33^ to infer base-wise conservation scores (SPH method, CONACC mode). Next, we calculated average phyloP (CONACC) scores for zCNEs present in each species, as well as exonic areas, and plotted conservation scores over the 25 zebrafish chromosomes as a heatmap using Circos ^29^. CNE synteny plotting for Figure 3 was created using the JCVI MCscan pipeline ^34^.

**Figure 3.**
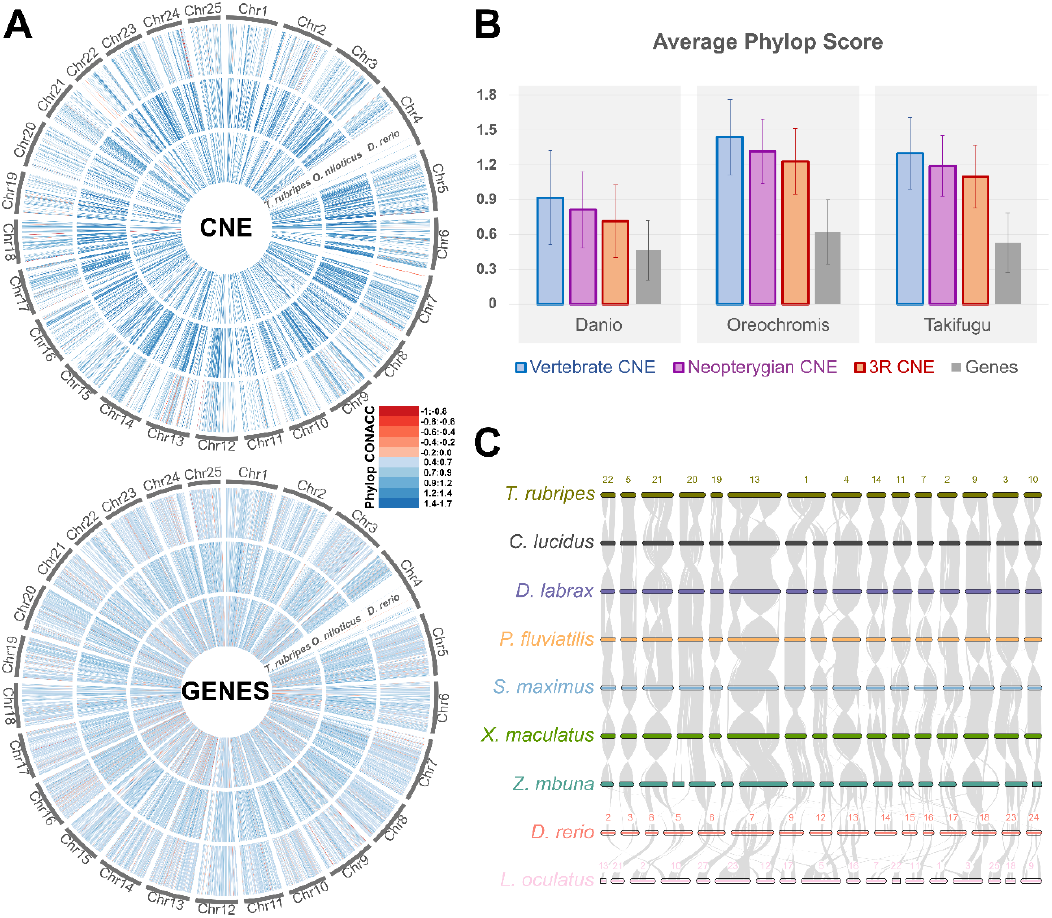
CNE sequence and synteny conservation. **A**. PhyloP (CONACC) Conservation scores for atCNEs and Genes in *D. rerio, O. niloticus* and *T. rubripes* mapped on *D. rerio* chromosomes. **B**. Average phyloP (CONACC) conservation scores and standard deviation for atCNEs in different categories and genes in *D. rerio, O. niloticus* and *T. rubripes*. **C**. Synteny of atCNEs across 8 teleost genomes and *L. oculatus*.

### Gene association and Ontology Enrichment

The association of CNEs with putative gene targets was carried out through an orthology-guided synteny-based approach. First, we extracted all genes that lie within 1Mb upstream and downstream of each CNE in each teleost species. A synteny score was attributed to each CNE - gene pair for zCNEs (Zebrafish as reference), fCNEs (Fugu as reference) or vCNEs (human as reference) which corresponds to the number of species in which the CNE is proximal to an orthologous gene, based on orthology information for all genes across species obtained by OrthoFinder2 v2.5.4. We then calculated a proximity rank for each target gene, which corresponds to the average position of a gene relative to a CNE, when all proximal genes are ordered in increasing distance (Figure 4A). Teleost CNEs were associated with candidate target genes with the highest synteny score (cutoff threshold of at least 5 species out of the 24 species scanned) and the lowest proximity rank (accepting all ties with the same syntenic score). For vCNE, we identified all target human genes within 1Mb of each CNE that are syntenic in human and either spotted gar or elephant shark at minimum. Associated atCNE targets in zebrafish or Fugu were then compared to vCNE targets, using Orthogroup information from OrthoFinder2. Gene targets of atCNE with orthologous loci found within 1Mb of vCNE were annotated as “ancestral targets” and gene targets without orthologous loci close to vCNE were annotated as “novel targets”. Furthermore, we used a bootstrap resampling approach to obtain support that the observed levels of association with ancestral targets were non-random. zCNEs or fCNEs were randomly associated with subsets of genes sampled from all zebrafish or Fugu genes which were compared to vCNE associated loci and the total number of ancestral or novel target associated CNE was counted. Using 10,000 bootstrap replicates with different random subsets, we calculated the average count, standard deviation and z score for ancestral or novel target associated elements and obtained strong support (z score <= 22) that associations were non-random. Gene ontology enrichment for GO biological process terms in different associated target gene sets was carried out through gProfiler ^35^.

**Figure 4.**
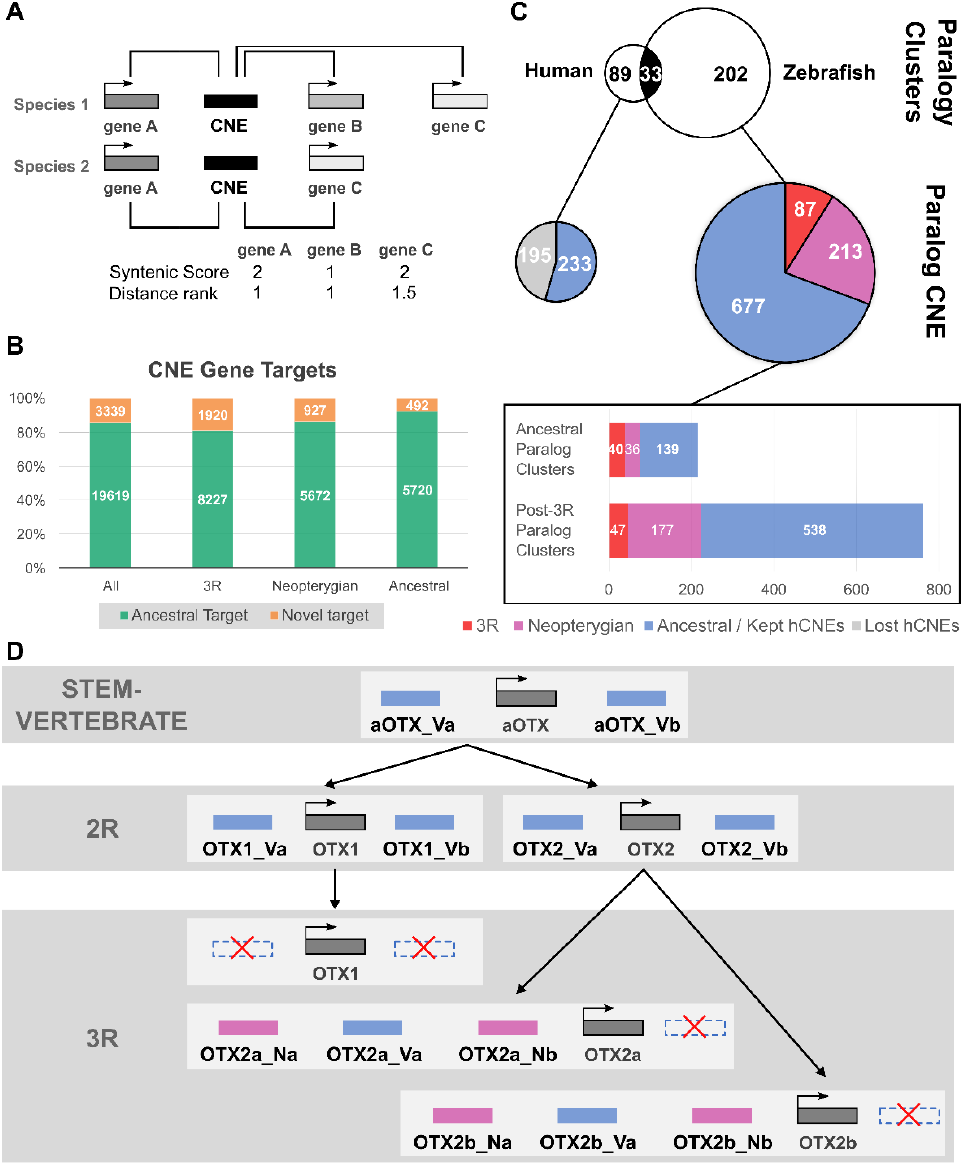
CNE-gene target association and paralog analysis. **A**. CNE were linked to putative target genes though orthology-guided synteny-based association, using the maximal syntenic score (number of species a CNE-gene pair is found within 1Mb) and minimal proximity rank (position of a gene to a CNE when all genes are ordered in increasing distance) for each CNE-target pair. **B**. Percentage (and counts within boxes) of ancestral (avCNE also proximal to orthologous locus in human) or novel gene targets for atCNEs in different categories. **C**. Paralogy cluster and paralog CNE counts for different categories in human / other vertebrates (left) or zebrafish / teleosts (right). Common paralogy clusters between the two groups are shared in black and common / ancestral atCNEs within paralogy clsuters are shown in blue. **D**. CNE gains, duplications and losses in the *OTX1* / *OTX2* paralogous loci in vertebrates and teleosts. The ancestral *OTX* locus in stem-vertebrate *aOTX* had two ancestral vertebrate CNEs (aOTX_Va & aOTX_Vb). Paralogous copies of these elements are found in human *OTX1* (OTX1_Va & OTX1_Vb) and *OTX2* (OTX2_Va & OTX2_Vb). In zebrafish, OTX1_Va, OTX1_Vb and OTX2_Vb were lost and the 3R WGD produced two paralogous copies of OTX2_Va (OTX2a_Va & OTX2b_Va). Two neopterygian gained CNEs in the *OTX2* locus (OTX2_Na & OTX2_Nb) also gave rise to two pairs of paralogous elements in the paralogous loci of *OTX2a* (OTX2a_Na & OTX2a_Nb) and *OTX2b* (OTX2b_Na & OTX2b_Nb).

### CNE Paralog Analysis

To identify CNE paralogs and clusters of paralogous CNE loci, we first carried out self-BLAST searches for zebrafish atCNEs and Fugu atCNEs to identify elements in each dataset with non-self hits. Gene paralog information for each species was then acquired from the Ensembl database and used to annotate putatively paralogous atCNE loci (as inferred from BLAST matches) based on the paralogy of their associated target genes. Gene paralog information was also used to infer the origin of paralogous atCNE loci prior or after the 3R WGD, based on the clade of origin of paralogous genes. In parallel, the same pipeline was applied to human vCNEs and inferred paralogous vCNEs were then compared to paralogous atCNEs to identify common paralogy clusters between the two groups that have orthologous loci associated with paralog CNEs in both groups.

### Motif Prediction and Transcription Factor Binding Site Enrichment

*De novo* motif prediction and transcription factor binding site enrichment analysis for different CNE sets were carried out using XSTREME online from the MEME suite. Enriched transcription factor binding sites (TFBS) were grouped to higher order categories based on similarity inferred from their consensus motifs and XSTREME motif clustering information. We also carried the same enrichment analysis on a subset of 10,932 random non-CNE zebrafish 24hpf embryonic enhancers downloaded from the EnhancerAtlas 2.0 database^36^ with comparable size distribution to CNE sets as a reference for motif and TFBS composition in non-conserved developmental enhancers.

## Results

### Teleost CNE identification and ancestral set reconstruction

For the *de-novo* identification of teleost CNEs, we first carried out whole-genome pairwise alignments in two pairs of focal reference species with high-quality annotations in selected positions of the phylogeny: 1) Zebrafish VS Mexican tetra and 2) Fugu VS gilthead sea bream. After filtering out coding and repetitive regions, this investigation resulted in a set of 298,743 conserved sequence chains between Zebrafish and Mexican tetra and 372,053 chains between Fugu and gilthead sea bream, yielding a total of 63,023 Zebrafish-Tetra CNEs (zCNE) and 39,532 Fugu-Seabream CNEs (fCNE) of at least 100bp with >=70% sequence identity.

We used the zCNE and fCNE datasets as reference CNE queries to scan the other teleost genomes via BLAST, while we also included the spotted gar genome as an outgroup to identify CNE gains and losses following the 3R WGD (Supplementary Tables 2-4 & 6). This approach allowed us to infer the ancestral teleost CNE set, by using presence/absence information of zCNEs and fCNEs in different species of our phylogeny. Taking advantage of the phylogenetic positions of our selected focal species, we carried out a bidirectional comparison, reviewing the presence of fCNE in the Zebrafish-Mexican tetra clade (Clade 1) and the presence of zCNE in the Fugu-gilthead sea bream clade (Clade 2), as described in more detail in the related material & methods section and summarized in Figure 1A. This search identified 7,358 fCNEs present in at least one species of Clade 1 and 22,596 zCNEs present in at least one species of Clade 2. Combining these sets produced a final set of 25,930 non-overlapping unique elements that consists the ensemble of CNEs inherited by at least two or more species in our phylogeny from the teleost common ancestor (atCNEs).

### Extensive CNE gain and loss during early teleost evolution

We next used the reconstructed atCNE set to review previously reported CNE loss in teleosts and to assess putative gains during early teleost evolution. We identified 6,585 atCNE that are also found in other vertebrates (Vertebrate atCNE), 6,934 atCNE that are shared only with spotted gar and no other vertebrates (Neopterygian atCNE) and 12,411 atCNE with no hit in vertebrates or spotted gar and are teleost specific (3R atCNE) (Figure 1A, Supplementary Table 7). We also used publicly available sets of CNEs from the ANCORA database, combining non-overlapping unique CNE shared by human-spotted gar and human-elephant ghost shark to obtain an ancestral Vertebrate CNE set (vCNEs). Searching for vCNEs in any of the teleost genomes in our phylogeny confirmed previously reported levels of loss of ancestral vCNEs in teleosts (69.5%) (Figure 1A, Supplementary Tables 5-6). A characteristic of vertebrate CNEs is their non-random distribution in the genome, with many clustering around developmental transcription factor genes. To assess if gained atCNEs follow a similar distribution pattern, we looked into the distribution of the three different categories of atCNEs in the zebrafish genome. After dividing the zebrafish genome in 500kb non-overlapping windows (hereinafter referred to as 500kb Zones), we counted total CNE content for each category in each zone (Supplementary Table 8). Of the total 2664 500kb zones, 1982 zones have atCNE presence, with 985 of these zones containing vertebrate atCNEs, 1179 zones containing neopterygian atCNEs and 1690 zones containing 3R atCNEs, while 682 zones have absence of atCNEs of any category (Figure1B).

### Evaluating atCNEs as Phylogenetic Markers

Taking advantage of our atCNE set, we assessed the potential of atCNEs as phylogenetic markers. We constructed two CNE based phylogenies, one with an extended dataset of 4,673 pan-teleost atCNEs present in all teleost species in the study but excluding spotted gar and one from a subset of the pan-teleost dataset with 2,668 elements also present in spotted gar. In parallel, we also constructed a protein-based phylogeny using universal single copy ortholog genes as a control for assessing resulting topologies from the CNE-based reconstructions. Use of the pan-teleost dataset of 4,673 atCNEs lead to the recovery of the expected topology, as supported by the gene-based tree (Supplementary Figure 3), with maximal bootstrap support for all clades (Figure 2A). In contrast, the dataset including spotted gar resulted in topological discrepancies in the relationships of *Lates calcarifer, Seriola dumerili* and *Scophthalmus maximus* and lower support in some of the clades (highlighted in red in Figure 2A).

### Variable Ancestral CNE loss across the Teleost Phylogeny

To characterize patterns of atCNE gain and loss in different teleosts, we mapped rates of loss for each atCNE category (Supplementary Table 9) as a percentage of all identified atCNEs, as presented in Figure 2B. A range of CNE loss is seen across the phylogeny, with different species having variable atCNE retention levels, while a pattern of incremental loss is seen for different atCNE categories (Figure2B). Vertebrate atCNEs have the lowest rates of loss ranging from 0.84% to 6.09%, with an average loss of 3.18% (standard deviation = 1.33%). In turn, neopterygian atCNEs show nearly two-fold loss in comparison, ranging from 1.08% to 10.53%, with an average loss of 5.31% (standard deviation = 2.56%). Finally, 3R atCNEs account for two thirds (average 68.1%) of total atCNE loss, with their loss ranging from 3.67% to 28.75% of all atCNEs, with an average of 17.82% (standard deviation = 6.78%).

### High Sequence and Synteny Conservation across teleost CNE

Vertebrate CNEs are characterized by high levels of sequence conservation, being on average more conserved than exonic areas^3^. To assess to what extent different categories of teleost CNEs follow this pattern of slow evolution, we carried out multiple whole genome alignment of the genomes included in our phylogeny and calculated phyloP conservation scores for atCNEs in Zebrafish, Fugu and Nile Tilapia (Supplementary Tables 10-13). We also assessed CNE synteny in chromosome level genomes included in our phylogeny, hypothesizing that the clustered distributional pattern of the majority of atCNEs around specific genomic regions suggested high synteny conservation. Teleost CNEs have significantly higher average phyloP scores compared to exonic sequences. What is more, vertebrate atCNEs have higher average phyloP scores than Neopterygian atCNEs, which in turn have higher phyloP scores than 3R atCNEs (Figure3B). CNE synteny is also largely conserved across teleosts, with larger syntenic areas shared among clade 2 species and notable rearrangements compared to zebrafish and spotted gar (Figure 3C, Supplementary Figure 4).

### Gene Association and Paralog Analysis Highlight Teleost CNE Evolution

Since CNEs have been shown to act as conserved developmental enhancers, we undertook a synteny based association approach to identify putative gene targets. (Figure 4A). Briefly, for each gene within 1Mb upstream or downstream of a zCNE or fCNE we attributed a synteny score (the number of species in which the CNE-gene pair is found) and a proximity rank (average position of each gene from a CNE in increasing order of distance). One or more targets (in case of tied candidates) with the highest syntenic score and the lowest distance rank were linked to each CNE (Gene A in Figure 4A example), using a minimum synteny score threshold of 5 species (Supplementary Tables 14-18).

Our gene association pipeline associated 20168 zCNEs with 4,444 zebrafish genes and an additional 2,790 fCNEs not corresponding to zCNEs with 1,146 Fugu genes. Comparing these sets to vCNE associated genes in human revealed that 85.46% of all atCNEs with targets were associated with a locus that has a human ortholog located within 1MB of a vCNE. We termed loci with human orthologs associated with vCNE “ancestral targets” and loci without human orthologs associated with vCNEs “novel targets” (Figure 4B). Looking at individual categories of atCNE with targets, 92.08% of vertebrate atCNEs, 85.95% of Neopterygian atCNEs and 81.08% of 3R atCNEs were associated with ancestral targets (Figure 4B). Gene ontology enrichment for ancestral atCNE targets showed high enrichment for genes involved in transcriptional regulation, the development of various organ systems and neuronal specification (Table 1, Supplementary Tables 19 & 20). Novel atCNE targets were mainly enriched for developmental processes already associated with ancestral target sets, but had significantly weaker enrichment support, while one process (regulated exocytosis) specifically enriched only in novel target genes (Table 1, Supplementary Tables 19 & 21).

**Table 1.**
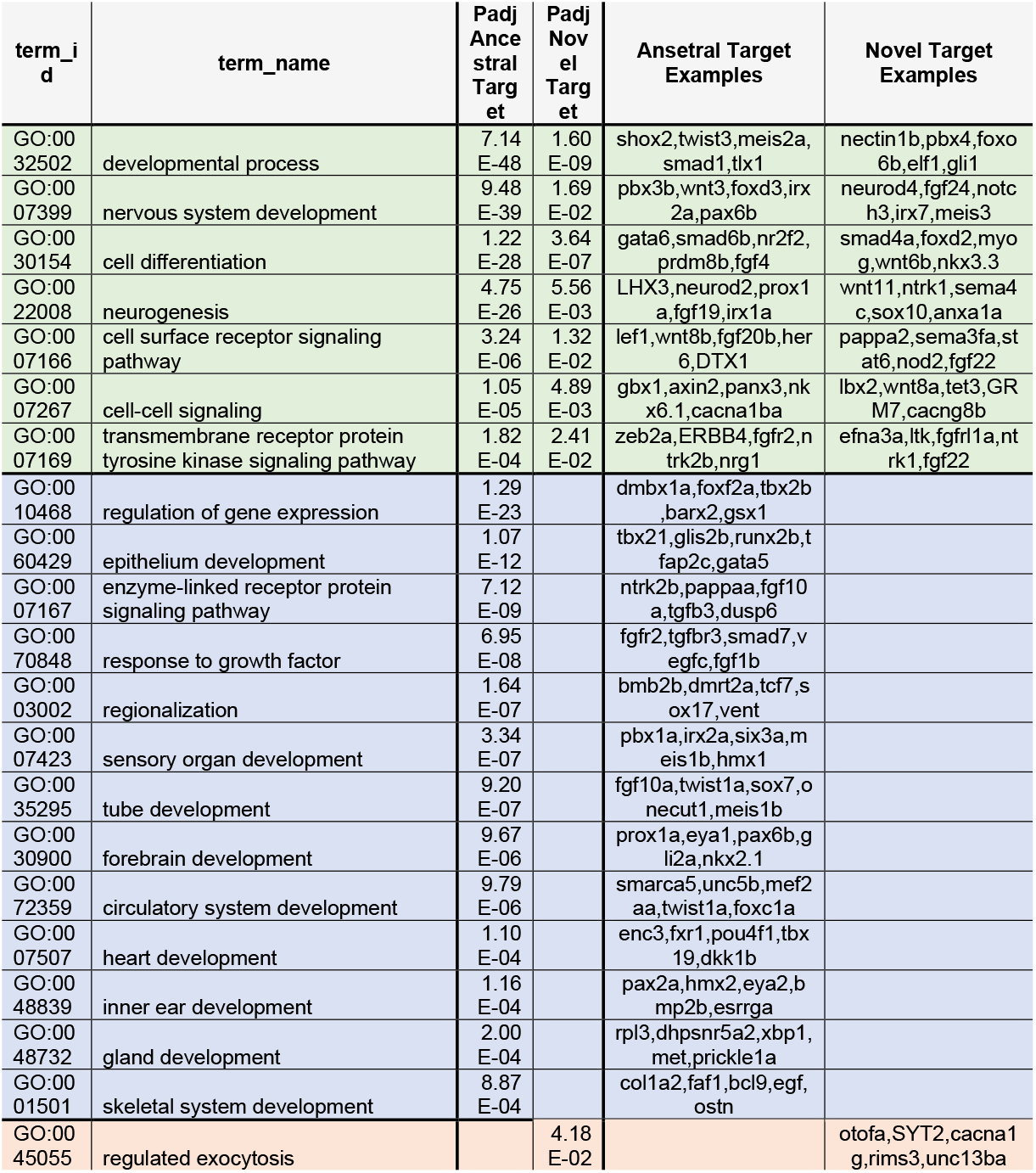
Gene Ontology Enrichment for ancestral and novel CNE gene target sets.

To identify WGD derived CNE copies, we combined self-BLAST searches of zebrafish and Fugu atCNEs with paralogy information from ENSEMBL, while applying the same methodology to human vCNEs in parallel, to compare vCNE and atCNE paralogous loci and probe ancestral CNE fate after the 2R and 3R WGD (Supplementary Tables 22-25). Our analysis identified 89 vCNE paralogy clusters in human and 202 atCNE paralogy clusters in zebrafish, with 33 common paralogy clusters shared by both groups (Figure 4C). Of the total 977 paralogous atCNEs, 69.3% are vertebrate atCNEs, 21.8% are Neopterygian atCNEs and 8.9% are 3R atCNEs, while 195 of 428 paralogous vCNEs were lost in teleosts (Figure 4C). A total of 215 atCNEs (22%) are located in ancestral (2R derived) paralog clusters, with the remaining 762 atCNEs (78%) found in 3R derived paralog clusters (Figure 4C). As an illustrative example of paralogous CNE evolution, gains and losses of paralogous CNEs in the *OTX1* / *OTX2* loci are presented in Figure 4D. Based on the structure of the locus in teleosts and other vertebrates, the ancestral *OTX* locus in stem-vertebrates (*aOTX*) contained two CNEs (aOTX_Va & aOTX_Vb), with each giving rise to a paralogous element in the paralogous loci of *OTX1* (OTX1_Va & OTX1_Vb) and *OTX2* (OTX2_Va & OTX2_Vb) after the 2R WGD. In teleosts, OTX1_Va, OTX1_Vb and OTX2_Vb were lost, while OTX2_Va is found in two paralogous copies (OTX2a_Va – OTX2b_Va) in the paralogous *OTX2a* and *OTX2b* loci. In parallel, two CNEs gained in the *OTX2* locus in the neopterygian ancestor (OTX2_Na & OTX2_Nb) gave rise to two pairs of paralogous elements in the *OTX2a* (OTX2a_Na & OTX2a_Nb) and *OTX2b* (OTX2b_Na & OTX2b_Nb) loci after the 3R WGD as well.

### Gained and Ancestral Teleost CNE Share Similar Motif Vocabularies

To assess if gained atCNEs may be divergent ancestral elements, we carried out motif and transcription factor binding site (TFBS) enrichment on different sets of atCNEs (Vertebrate, Neopterygian, 3R) and vCNEs (Lost and Kept in Teleosts) (Supplementary Tables 26-32). In parallel, as a control, we also carried out the same enrichment analysis on 10,932 random non-CNE zebrafish enhancers with similar size distribution to our CNE sets (Supplementary Table 33). All CNE categories, but not non-CNE enhancers, were found to share a common TFBS vocabulary, dominated by the homeobox (40-60% of CNE), FOX (35%-61%), EST-related / C2H2 zinc finger (28%-47%) and bHLH (15%-33%) classes (Figure 5). *De novo* motif discovery also found an enrichment for the “TAATTA” homeodomain binding motif, centrally positioned in 18%-24% of all CNEs (Figure 5). No shared enriched motifs were found between any atCNE and vCNE combinations, but 19.7%-53.3% of atCNEs contained Prdm5 sites, which were not enriched in vCNEs (Figure 5). Additionally, 14.8% of 3R atCNEs contained PAX2 sites, which were not specifically enriched in any other CNE category.

**Figure 5.**
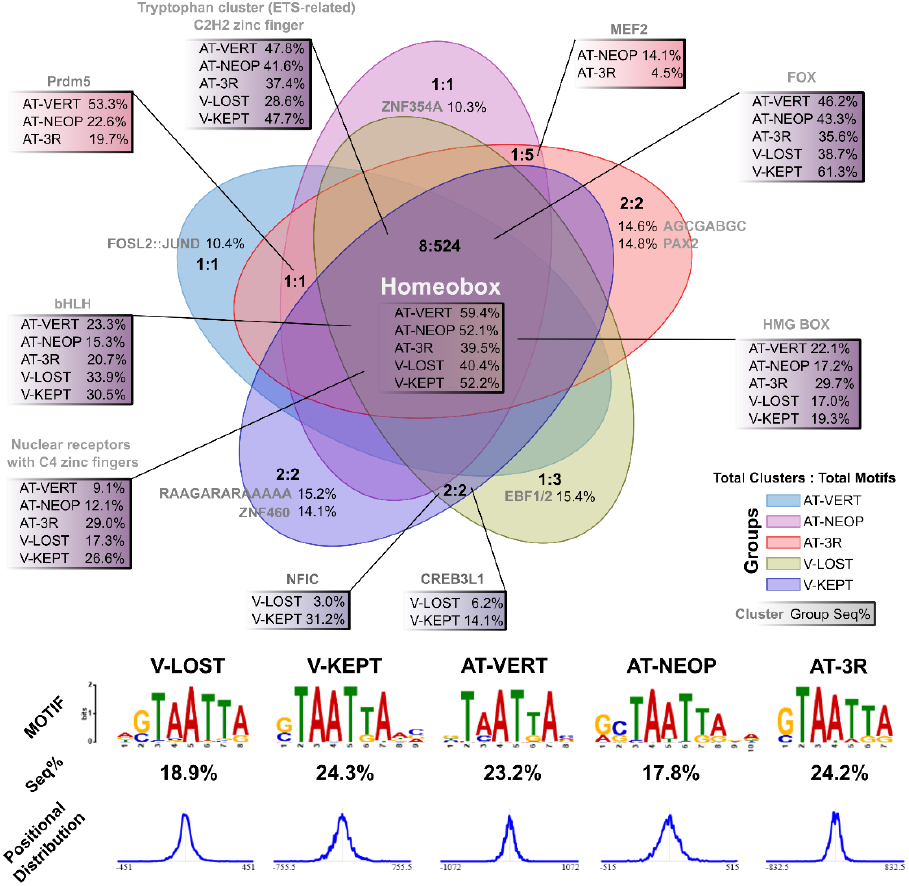
Motif Discovery and Transcription Factor Binding Site Enrichment. **TOP**. Overlap of enriched TFBS categories among combinations of atCNEs (AT-VERT : Vertebate - Blue, AT-NEOP Neopterygian - Purple, AT-GAIN : 3R - Red) and avCNEs (V-KEPT : Kept in Teleosts - Dark Purple, V-LOST : Lost in Teleosts - Yellow). Enrichment overlaps are displayed in the format “Total Categories: Total Enriched Motifs” in each overlap area. The percentage of sequences of each CNE group with motifs in representative categories (c1, c2…) are shown. **BOTTOM**. Percentage of sequences in each CNE group that contain the most common shared *de novo* predicted motif (consensus TAATTA) and its positional distribution.

## Discussion

This work presents the first attempt to catalogue the previously undocumented variety of neopterygian and teleost CNEs by using a teleost-centric identification protocol. This approach is able to achieve high levels of CNE identification with low to moderate time and resource requirements, combining thorough pairwise whole genome alignments for strategically selected focal species with faster BLAST scanning for a wider range of teleosts. This offers an attractive methodology for fast CNE searches in large genomic datasets, representing an efficient way for answering questions of ancestral CNE gains and losses. Our work also offers support on the value of CNEs as a tool for robust phylogenomic reconstruction of teleost relationships, drawing attention on the importance of appropriate outgroup selection, as in the case of spotted gar shared elements.

### Properties of Teleost CNE

Core aspects of the identity and evolution of teleost CNE are highlighted here. Importantly, our work shows that the loss of ancestral CNEs in early teleosts was accompanied by extensive gain of new elements. While a large number of new CNEs (35.8%) were gained in the neopterygian ancestor, the majority of new elements (64.2%) were acquired after the split with spotted gar, following the 3R WGD. The study of the genomic distribution of ancestral and gained elements in zebrafish also provides further insight into how these elements were acquired. The majority of genomic zones containing vertebrate atCNEs (89.2%) also contain gained CNEs, with 80.4% of neopterygian atCNEs and 84.7% of 3R atCNEs found in ancestral vertebrate zones. Respectively, novel zones without vertebrate atCNEs have much lower total CNE content, with novel neopterygian zones (40.2% of all zones with neopterygian atCNEs) containing only 19.6% of all neopterygian atCNEs and novel 3R zones (30.9% of all zones with 3R atCNEs) containing only 15.3% of all 3R atCNEs. In addition, a quarter of the zebrafish genome (25.6%, *i.e*. 682 / 2664 zones) is devoid of CNEs, supporting that all categories of teleost CNEs (whether ancestral or gained) exhibit non-random clustered distribution, which is distinct from the more homogeneous distribution of genic regions (Figure 3A).

Overall, key properties of teleost CNEs are elucidated here: 1) they are distributed over the similar overlapping genomic regions, regardless of their time of origin, 2) they show high sequence and synteny conservation over large evolutionary distances, comparable to CNEs in other vertebrates ^3,37^, 3) they are syntenically associated with loci involved in development related processes and transcription factor genes and 4) they are enriched for development related transcription factor binding sites, with many harboring homeodomain binding motifs centrally in their sequence. The latter characteristic is in line with previous reports of AT enrichment in the conservation core of vertebrate CNEs, suggested to emerge from an overrepresentation of homeodomain binding motifs^38,39^.

### The Evolutionary Dynamics that shaped the teleost CNE Repertoire

The massive loss of ancestral CNEs in teleosts has been considered a mystery since early studies of vertebrate CNEs, mainly because the expected impact of the 3R WGD would be an addition of new paralogous CNEs to existing elements incrementally. Consequently, finding that novel CNEs gained in stem-teleosts have highly similar characteristics as ancestral vertebrate elements is pivotal for advancing our understanding of CNE evolution. While possibly counter-intuitive at first, this re-establishment of the CNE repertoire following the 3R WGD is comparable to the fixation of ancestral CNEs in the jawed vertebrate common ancestor. With only few hundreds of CNEs shared with lamprey and only tens of elements recognizable in amphioxus, the vast majority of jawed vertebrate CNEs were fixed after the second WGD in jawed vertebrates^40^. CNE associated targets, as well as experimentally confirmed regulated targets, include loci involved in the development of the central and peripheral nervous systems, as well as heart and muscle during the specification of these tissues at the phylotypic stage^41^. However, most of the innovations related with these cell populations and their associated gene regulatory networks predate the split of jawed vertebrates and are shared with cyclostomes. Similarly, we observed that the majority of teleost gained CNEs are also associated with these ancestral targets involved in developmental regulation at the phylotypic stage. This is in contrast with younger clade specific CNE gains in mammals, which have been associated with novel target loci, many of which are involved in protein-binding ^42,43^. Similarly, previous work showed that older vertebrate CNEs are implicated in transcriptional and developmental regulation, while younger mammalian elements are associated with genes involved in post-transcription modification^44^. Finally, gained (either Neopterygian or 3R) teleost CNE also share a highly similar motif and binding site composition as vertebrate CNE, supporting they are also regulated by similar upstream networks. Consequently, we hypothesize that the majority of gained CNEs in teleosts represent a reconfiguration of the ancestral vertebrate CNE repertoire linked to the regulation of the same key ancestral targets, instead of components of a new network regulating teleost specific processes.

### CNE Gain by Sequence Divergence and De Novo Emergence

While atCNEs of different origin categories have largely comparable characteristics, there are differentiating features that are detected when comparing gained (Neopterygian or 3R) and ancestral (vertebrate) teleost CNE. When considered in decreasing age of conserved sequence fixation (Vertebrate > Neopterygian > 3R), older atCNEs show higher average conservation, have lower loss rates across the phylogeny and are associated with more ancestral target genes compared to newer elements. These differences may originate from a small number of neopterygian and 3R atCNEs that are located within novel genomic regions without vertebrate atCNEs and which are associated with novel teleost specific targets (Figure 6A). Based on this deviating profile compared to other atCNEs, it can be suggested that these novel-target associated elements are cases of *de novo* CNE emergence, as part of teleost specific regulatory networks. We therefore hypothesize that the ensemble of the CNEs gained in the stem-teleost lineage derived from two complementary scenarios of evolutionary origin (Figure 6B). The homogeneous profile of the majority of atCNEs is compatible with a scenario of CNE gain through extreme sequence divergence, in which ancestral vertebrate elements deviate in primary sequence, while retaining ancestral TFBS content and remaining associated with ancestral targets. This must have then been coupled with *de novo* gains, as supported by both the emergence of novel-target teleost specific elements and the evolution of Neopterygian CNEs around ancestral targets prior to the extensive CNE loss in teleosts. The analysis of paralogous CNE evolution further illustrates this, with more than 1 in 5 paralogous atCNEs having neopterygian origin (21.8%), as in the case of the *OTX1* / *OTX2* loci where gain of neopterygian paralogous atCNEs was coupled with loss of ancestral vertebrate paralogous CNEs.

**Figure 6.**
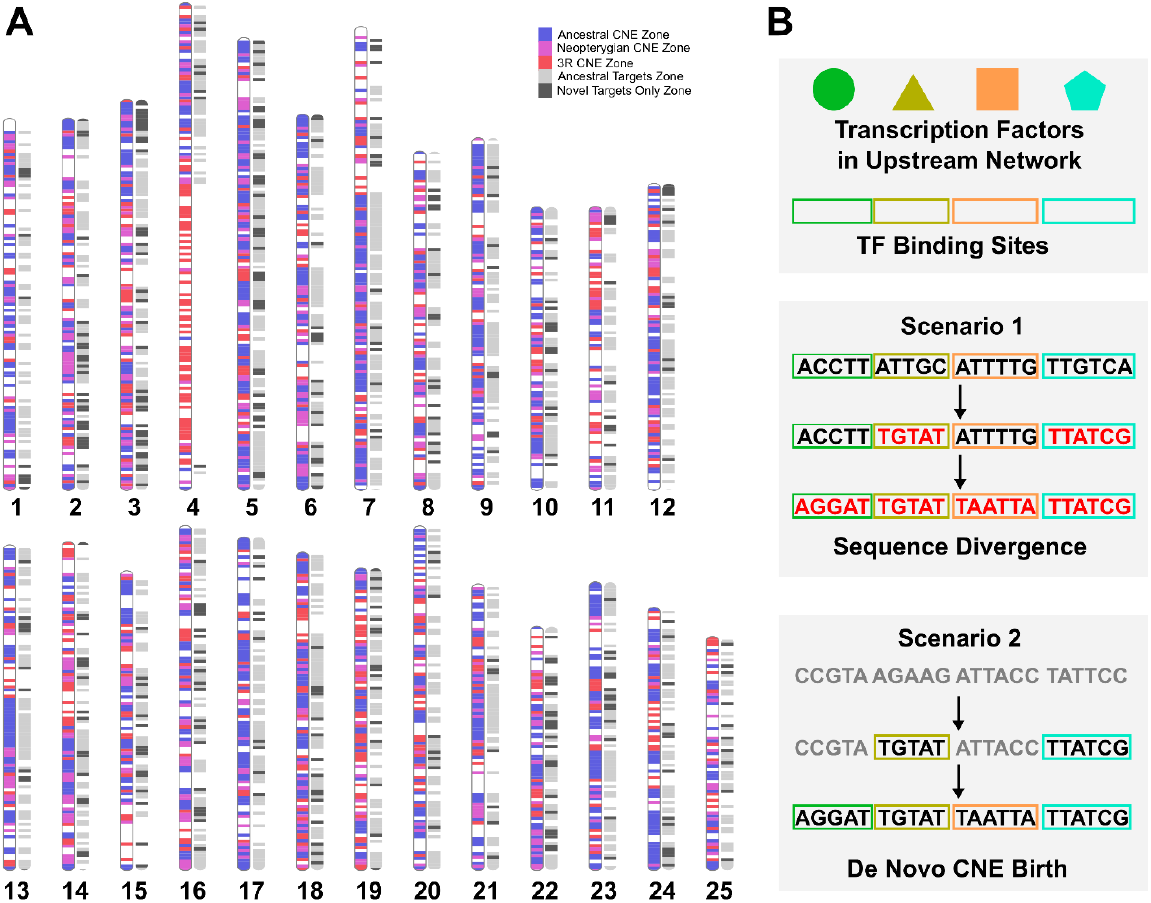
Teleost CNE-target localization and models of CNE gain. **A**. Localization of atCNEs and associated target genes in zebrafish chromosomes. CNE zones are plotted in blue if they contain vertebrate atCNEs, in purple if they contain neopterygian atCNEs and do not contain vertebrate atCNEs, in red if they contain 3R atCNEs and do not contain any older (vertebrate or neopterygian) atCNEs. Gene zones are plotted in light grey if they contain ancestral targets or in dark grey if they contain only novel targets and do not contain ancestral targets. **B**. Two hypothesized scenarios of atCNE gain: scenario 1) Extreme sequence divergence of ancestral sequence renders CNE unrecognizable, while TFBS are preserved and scenario 2) CNE gain through *de novo* emergence.

## Author contributions

Study conceptualization and design: V.P. & T.M., Data Analysis: E.I. & V.P., Interpretation: E.I., V.P., C.S.T. & T.M., Project Supervision C.S.T. & T.M., Manuscript Preparation: E.I., V.P., C.S.T. & T.M.

## Acknowledgements

This research was supported through computational resources provided by IMBBC of the HCMR. Funding for establishing the IMBBC HPC has been received by the MARBIGEN (EU Regpot) project, LifeWatchGreece RI and the CMBR (Centre for the study and sustainable exploitation of Marine Biological Resources) RI. The authors want to thank Dr Pavlos Pavlidis for support on the initial setup of the project and the co-supervision of the MSc thesis of E.I.

## Data Availability

Publicly available genomic data used for the analyses in this work are listed in supplementary data. Data generated as part of this study (including CNE genomic coordinates) are provided through the supplementary data document and supplementary tables 1-33 in supplementary files 1-5, with table description included in the supplementary data document. New code generated and used in this project will be made publicly available through https://github.com/genomenerds/CNE_analysis.

