## Supplementary Data for "Extensive Loss and Gain of Conserved Non-Coding Elements during Early Teleost Evolution"

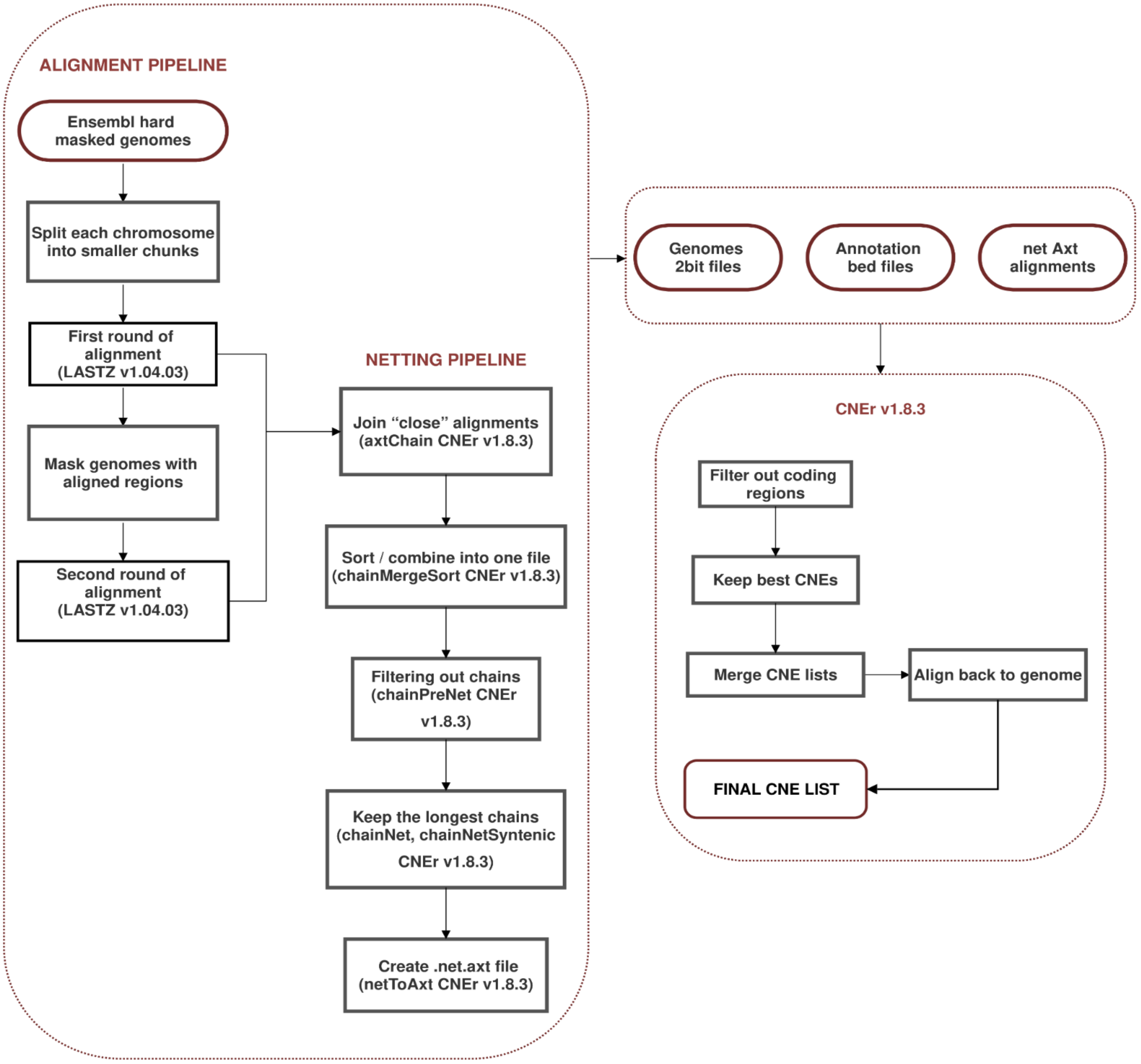


**Supplementary Figure 1. Alignment pipeline**. Two rounds of pairwise whole-genome alignments using masked genomes. **Netting pipeline**. Join close alignments and form chains, then select the longest chains. **CNEr v1.8.3.** CNE Identification pipeline using genomes in 2bit format, annotation bed files and the chained alignments as input. The output of this pipeline is a list with all identified CNEs.

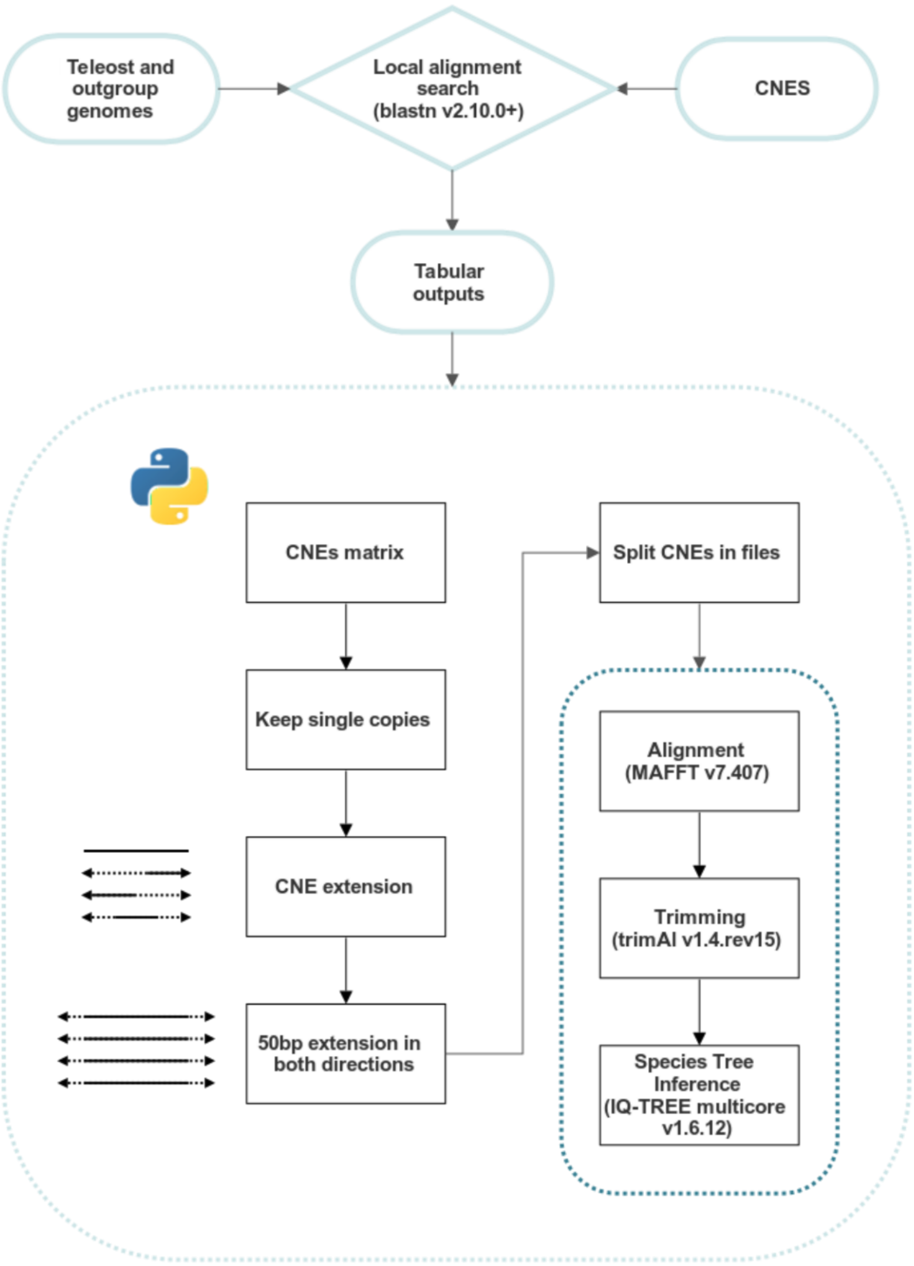


**Supplementary Figure 2.** CNE processing pipeline and CNE-based tree construction using single copy CNE orthologs.

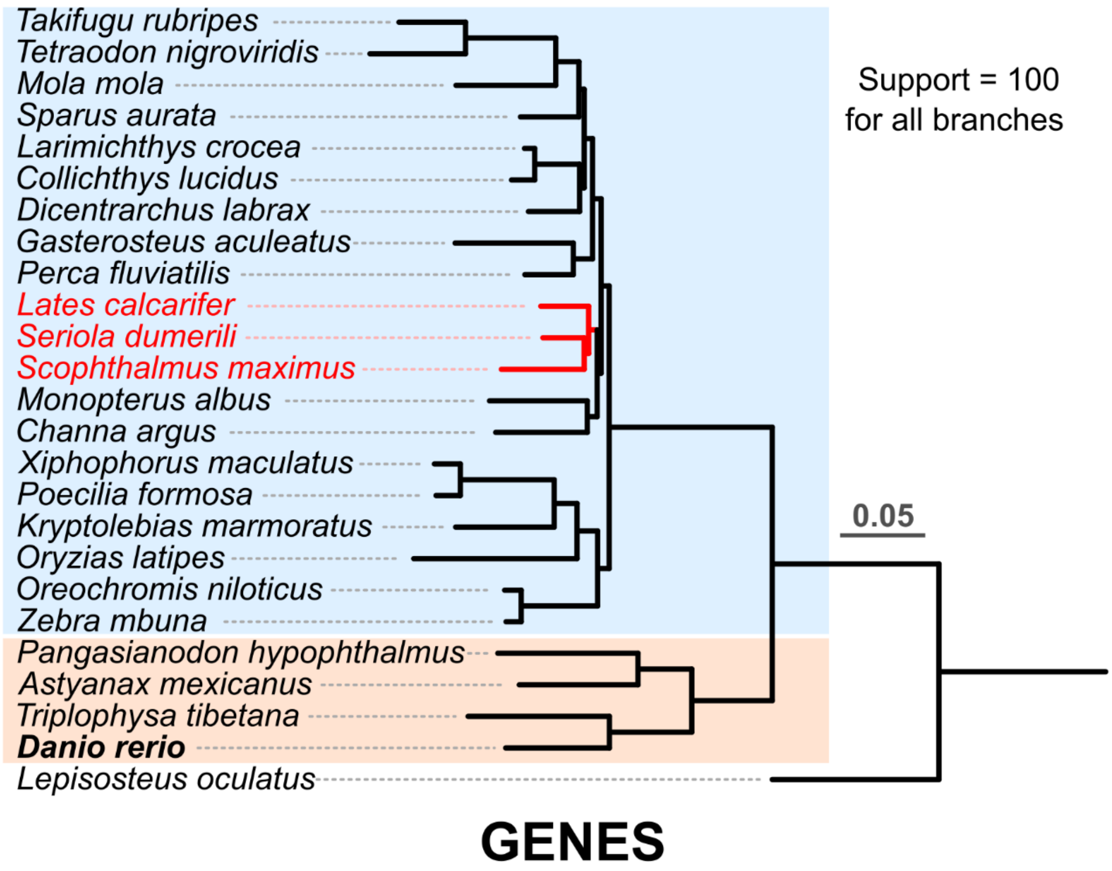


**Supplementary Figure 3.** Phylogenetic reconstruction of the relationships of the 24 teleost species included in the study, using single copy universal ortholog genes (identified via OrthoFinder – see materials and methods for details). The two basally diverged clades used to infer ancestral teleost CNEs for main Figure 2A are highlighted with orange (clade 1) and blue (clade 2). Branches with species highlighted in red show topological discrepancies in the phylogeny built only with spotted gar shared atCNEs (main Figure 2A).

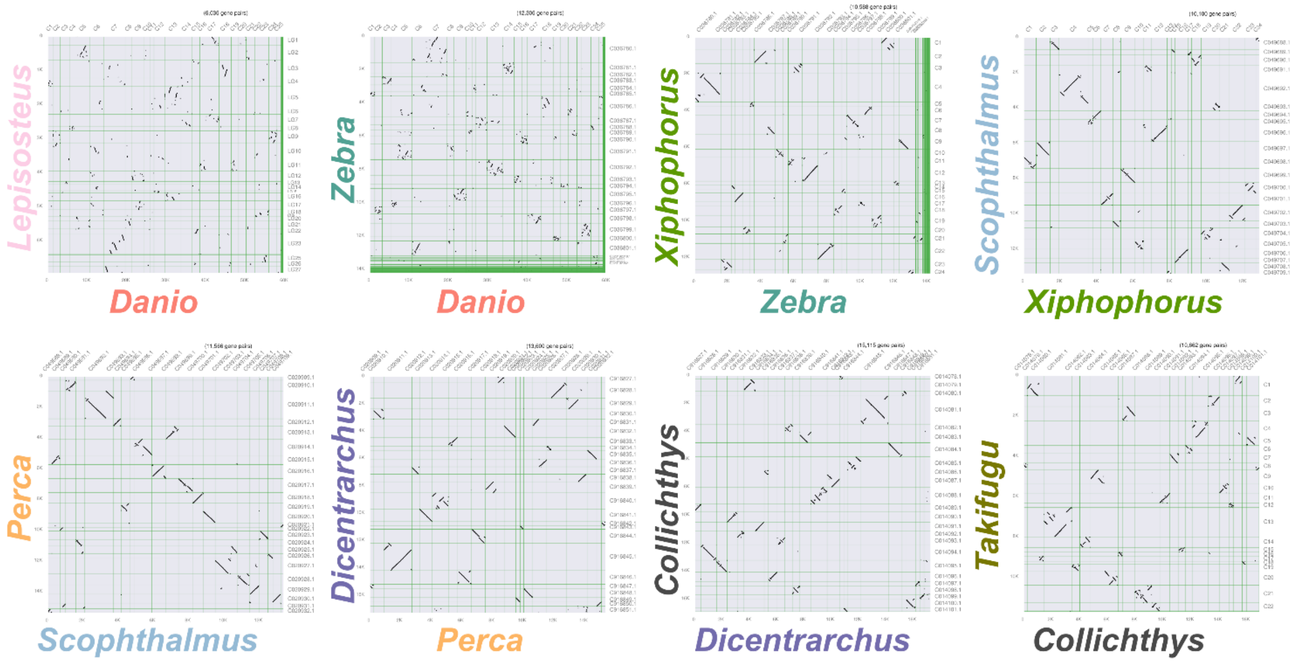


**Supplementary Figure 4.** Syntenic dotplots for species used for synteny conservation analysis for main Figure 3C.

**Supplementary Table 1. Genome versions used in this study.**

| Species | Genome | Source |
|---|---|---|
| Astyanax_mexicanus | Astyanax_mexicanus-2.0 | ENSEMBL |
| Channa_argus | GCA_004786185.1_ASM478618v1 | NCBI RefSeq |
| Collichthys_lucidus | GCA_004119915.2_ASM411991v2 | NCBI RefSeq |
| Danio_rerio | GRCz11 | ENSEMBL |
| Dicentrarchus_labrax | seabass_V1.0 | ENSEMBL |
| Gasterosteus_aculeatus | BROADS1 | ENSEMBL |
| Kryptolebias_marmoratus | ASM164957v1 | ENSEMBL |
| Larimichthys_crocea | L_crocea_2.0 | ENSEMBL |
| Lates_calcarifer | ASB_HGAPassembly_v1 | ENSEMBL |
| Mola_mola | ASM169857v1 | ENSEMBL |
| Monopterus_albus | M_albus_1.0 | ENSEMBL |
| Oreochromis_niloticus | O_niloticus_UMD_NMBU | ENSEMBL |
| Oryzias_latipes | ASM223467v1 | ENSEMBL |
| Pangasianodon_hypophthalmus | GCA_009078355.1_GENO_Phyp_1.0 | NCBI RefSeq |
| Perca_fluviatilis | GCA_010015445.1_GENO_Pfluv_1.0 | NCBI RefSeq |
| Poecilia_formosa | PoeFor_5.1.2 | ENSEMBL |

| | | |
|---|---|---|
| Scophthalmus_maximus | GCF_013347765.1_ASM1334776v1 | NCBI RefSeq |
| Seriola_dumerili | Sdu_1.0 | ENSEMBL |
| Takifugu_rubripes | fTakRub1.2 | ENSEMBL |
| Tetraodon_nigroviridis | TETRAODON8 | ENSEMBL |
| Triplophysa_tibetana | GCA_008369825.1_ASM836982v1 | NCBI RefSeq |
| Xiphophorus_maculatus | X_maculatus-5.0-male | ENSEMBL |
| Zebra_mbuna | GCF_000238955.4_M_zebra_UMD2a | NCBI RefSeq |
| Lepisosteus_oculatus | LepOcu1 | ENSEMBL |
| Homo_sapiens | GRCh38.p13 | NCBI RefSeq |
| Gallus_gallus | GRCg6a | NCBI RefSeq |
| Callorhinchus_milii | Callorhinchus_milii-6.1.3 | NCBI RefSeq |
| Anolis_carolinensis | anoCar2 | NCBI RefSeq |
| Xenopus_tropicalis | UCB_Xtro_10.0 | NCBI RefSeq |

**Supplementary Table 2 (t2_zCNE_blast in Supplementary File 1).** Coordinates of all zCNEs based on BLAST searches in different teleost species.

**Supplementary Table 3 (t3_fCNE_blast in Supplementary File 1).** Coordinates of all fCNEs based on BLAST searches in different teleost species.

**Supplementary Table 4 (t4_fCNE_NONzCNE_blast in Supplementary File 1).** Coordinates of fCNEs not overlapping zCNEs based on BLAST searches in different teleost species.

**Supplementary Table 5 (t5_vCNE_blast in Supplementary File 1).** Coordinates of vCNEs (Human sequences of vertebrate ancestral CNEs from ancora db shared with spotted gar or elephant shark – see materials and methods) based on BLAST searches in different teleost species.

**Supplementary Table 6 (t6_blast_counts in Supplementary File 1).** Count of total zCNEs, fCNEs, fCNEs not overlapping zCNEs and vCNEs identified in each teleost species via BLAST.

**Supplementary Table 7 (t7_atCNE_unmerged in Supplementary File 2).** Vertebrate, neopterygian and 3R atCNEs (unmerged coordinates from CNEr).

**Supplementary Table 8 (t8_danio_genome_CNE_counts in Supplementary File 2).** Total count of atCNE in each category across 500kb zones in *D. rerio* chromosomes.

**Supplementary Table 9 (t9_atCNE_presence in Supplementary File 2).** Counts of atCNEs from different categories present in each teleost species.

**Supplementary Table 10 (t10_phyloP in Supplementary File 1).** phyloP (CONACC) score for atCNEs in *D. rerio*, *O. nilocitus* and *T. rubripes.*

**Supplementary Table 11 (t11_danio_coding_phylop Supplementary File 2).** phyloP (CONACC) score for coding regions in *D. rerio.*

**Supplementary Table 12 (t12_oreochromis_coding_phylop in Supplementary File 2).** phyloP (CONACC) score for coding regions in *O. nilocitus.*

**Supplementary Table 13 (t13_takifugu_coding_phylop in Supplementary File 2).** phyloP (CONACC) score for coding regions in *T. rubripes.*

**Supplementary Table 14 (t14_atCNE_targets in Supplementary File 3).** Target genes associated with atCNEs.

**Supplementary Table 15 (t15_vCNE_1Mb_syntenic in Supplementary File 3).** Genes syntenically linked found within 1Mb of vCNEs.

**Supplementary Table 16 (t16_target_gene_orthology in Supplementary File 3).** Orthology of atCNE target genes and genes found within 1Mb of vCNEs.

**Supplementary Table 17 (t17_gene_assoc_stats in Supplementary File 3).** Gene association stats for different categories of atCNEs, associated with ancestral and novel gene targets (see materials and methods for more details).

**Supplementary Table 18 (t18_gene_assoc_BOOTSTRAPING in Supplementary File 3).** Bootstrap resampling analysis results for randomly sampled atCNE gene targets.

**Supplementary Table 19 (t19_assoc_gene_gprofiler in Supplementary File 3).** GO enrichment (gProfiler) analysis results for ancestral and novel atCNE gene targets (GO biological terms).

**Supplementary Table 20 (t20_ancestral_Goterms in Supplementary File 3).** GO biological term annotations for ancestral atCNE gene targets.

**Supplementary Table 21 (t21_novel_Goterms in Supplementary File 3).** GO term annotations for novel atCNE gene targets.

**Supplementary Table 22 (t22_teleost_paralogy in Supplementary File 4).** Paralogy clustering analysis results for teleost atCNEs. atCNEs are grouped by associated locus (one per line) and paralogous loci are organized in paralogy groups (column A). Paralogous loci inferred to be a product of the 3R WGD are indicated in the “teleost_flag” cell (column J).

**Supplementary Table 23 (t23_human_paralogy in Supplementary File 4).** Paralogy clustering analysis results for human vCNEs. vCNEs are grouped by associated locus (one per line) and paralogous loci are organized in paralogy groups (column A).

**Supplementary Table 24 (t24_paralog_group_orthology in Supplementary File 4).** Orthology of paralogous teleost and human CNE containing loci (one teleost and one human locus per line).

**Supplementary Table 25 (t25_CNE_paralog_stats in Supplementary File 4).** CNE paralogy analysis summary stats.

**Supplementary Table 26 (t26_motif_enrichment_all_clusts in Supplementary File 5).** Clusters of enriched motifs found in different atCNE and vCNE categories. For each cluster the most common motif (motif found in the highest number of sequences) is indicated.

**Supplementary Table 27 (t27_motif_enrichment_compare in Supplementary File 5).** Stats and comparisons of clusters shared by different groups of atCNEs and vCNEs.

**Supplementary Table 28 (t28_vCNE_KEPT_XSTREME in Supplementary File 5).** Motif prediction and enrichment analysis results by XSTREME (see materials and methods for more details) for vCNEs kept in teleosts.

**Supplementary Table 29 (t29_vCNE_LOST_XSTREME Supplementary File 5).** Motif prediction and enrichment analysis results by XSTREME (see materials and methods for more details) for vCNEs lost in teleosts.

**Supplementary Table 30 (t30_atCNE_VERT_XSTREME in Supplementary File 5).** Motif prediction and enrichment analysis results by XSTREME (see materials and methods for more details) for vertebrate atCNEs.

**Supplementary Table 31 (t31_atCNE_NEOP_XSTREME in Supplementary File 5).** Motif prediction and enrichment analysis results by XSTREME (see materials and methods for more details) for neopterygian atCNEs.

**Supplementary Table 32 (t32_atCNE_3R_XSTREME in Supplementary File 5).** Motif prediction and enrichment analysis results by XSTREME (see materials and methods for more details) for 3R atCNEs.

**Supplementary Table 33 (t33_random_24hpf_danio_enhancer in Supplementary File 5).** Motif prediction and enrichment analysis results by XSTREME (see materials and methods for more details) for random 24hpf *D. rerio* enhancers (control - see materials and methods for more details).
